# Discrimination of Isomeric Glycans via High-sensitive Engineered Nanopore

**DOI:** 10.1101/2023.12.28.573513

**Authors:** Guangda Yao, Yinping Tian, Wenjun Ke, Jie Fang, Shengzhou Ma, Tiehai Li, Xi Cheng, Bingqing Xia, Liuqing Wen, Zhaobing Gao

**Author notes:** Corresponding author. (Z.G.), (L.W.), (B.X.), (X.C.). Mengyao Ma, Yinping Tian, and Wenjun Ke contributed equally to this work.

## Abstract

Glycan isomers are the basis for forming the spatial structure of glycans, which poses great challenges to glycan structural analysis and hinders the study of the relationship between structure and function. An economical and convenient analytical method with high resolution is still in demand. Here, we designed an engineered α-hemolysin nanopore with double-mutants (α-HL M113R/T115A) to achieve sensing of glycans with different glycosidic linkages. By extracting three parameters to depict a 3D scatter plot, four glycans with different degrees of polymerization and glycosidic bonds could be distinguished. Through molecular dynamics (MD), we further elucidated the motion trajectories of two different glycosidic bonds in the pore region. In addition, the sensing ability was also applied in the direct identification of glycans in mixture systems. The obtained nanopore data permitted us to achieve a separation accuracy of over 90% independent of the assistance of machine learning algorithms. Designing biological nanopores with high discrimination and sensitivity for glycans with different lineages paves the way for nanopore glycan profiling and potentially sequencing.

## INTRODUCTION

Glycan, composed of multiple monosaccharide units, constitutes the carbohydrate portions of glycoconjugates like proteoglycans ^1^, glycolipids ^2^, and glycoRNAs ^3^. Glycan plays diverse and essential roles in numerous biological systems by mediating specific molecular recognition events critical for embryonic development, cell growth, immune response, and disease progression ^4^ and is related to drug development ^5^ and disease diagnosis ^6^. Multiple glycoconjugates comprised of repeating Gal-GlcNAc units are likely to be potential biomarkers for cancer ^7^, virus infections ^8^, and quality determinants for human milk oligosaccharides (HMOs) ^9^. Isomerism (including differences in glycosidic linkages and anomeric configurations, etc.), a ubiquitous phenomenon in glycobiology, is necessary to maintain the binding specificity and relevant functions of glycoconjugates ^10^. The development and progression of many diseases are related to abnormalities of glycan isomerism ^11-13^. According to the anomeric configuration and position of the related carbon atom, glycosidic linkages can be divided into α type (α-1,1, α-1,2, α-1,3, α-1,4, α-1,5, α-1,6) and β type (β-1,2, β-1,3, β-1,4, β-1,6), which constitute different saccharide units and glycans with various biological functions ^14^. Accordingly, the Gal-GlcNAc units can be classified into Galβ1,4GlcNAc (LacNAc) and Galβ1,3GlcNAc (lacto-N-biose, LNB) with distinct biological functions as well ^15^. It is the considerable structure diversity of the glycan that confers glycoconjugates diverse biological functions ^16, 17^. Therefore, characterizing glycan structure may help to understand the significant roles glycan plays in physiology and pathology ^16^.

However, structure analysis of glycan still faces a great challenge due to the complexity and heterogeneity of glycan structure and the limitations of current detection technology ^18, 19^. Besides monosaccharide diversity, the isomerism caused by different anomeric configurations and glycosidic linkages also posed a great challenge to glycan identification. At present, nuclear magnetic resonance (NMR) spectroscopy including two/three-dimensional NMR (e.g., 2D-COSY, HMQC, HMBC, TOCSY, NOESY) ^20, 21^, mass spectrometry (MS) and MS-based technologies (e.g., LC-MS/MS, IM-MS, MS-IRMPD) ^22, 23^ are still the most widely used methods for glycan analysis. Both of them can be used to discriminate α/β anomers and to conclude possible sequences of glycans but are limited by glycan availability. Glycan microarray is powerful in exploring glycan-glycan binding protein (GBP) interactions but less useful in individual glycan molecule analysis ^24^. Besides, recognition tunnelling (RT) shows potential in classifying anomers and epimers by current alteration ^25^. However, none of these methods can provide full composition and structural information of glycans owing to detection principle limitations. For instance, both it is challenging for NMR/MS to characterize types of glycosidic bonds in glycans, which is quite necessary for glycan analysis. Many of the above methods are time-consuming and involve a series of complex procedures ^23^. Hence, a more comprehensive and efficient technology is urgently needed.

Nanopore technology, which emerged as a powerful ultrasensitive analytical tool with portability and lower costs, has been used in the analysis of DNA, RNA, protein, metabolites of body fluids, etc. ^26-28^. Compared with solid-state nanopores, biological nanopores have a more precise and reproducible structure and consist of amino acids which play a pivotal role in pore-analyte interaction, thus having more applications in bioanalyte analysis ^29, 30^. During a typical detection process using a nanopore, the analytes are driven through the nanopore from one side to the other side of a chamber separated by bilayer lipids under an applied voltage, inducing ionic current alteration while interacting with the sensing region, from which the structure and composition may be elucidated ^26^. During the last two decades, nanopore sequencing in DNA has made great progress, showing its potential in biopolymer analysis ^26, 28, 31^. Recently, more and more researchers began to explore the feasibility of analyzing glycans with nanopores. For instance, Zhang S et.al reported the identification of common monosaccharides using modified MspA ^32^. Aerolysin was used for the differentiation of glycosaminoglycans and the identification of glycan stereoisomers with chemical tags ^33, 34^. α-Hemolysin (α-HL), the first biological nanopore used in nucleotide analysis, with a long sensing region, may have the potential to discriminate minor differences in glycans as well ^35^. We have previously mapped the acetamido and carboxyl group on glycans at monosaccharide, disaccharide, and trisaccharide levels with engineered α-HL ^36^. Here, we have achieved discrimination between glycans with distinct glycosidic bonds with high resolution using another engineered α-HL nanopore and further elucidate the underlining mechanisms with molecular dynamics (MD) simulations.

## RESULTS AND DISCUSSION

### Detection of glycans with distinct glycosidic bonds

The engineered α-hemolysin nanopore with a single R mutation at its narrowest region was reported previously and was utilized for the first time to map the acetamido groups by the features of blockage events at the monosaccharide, disaccharide, and trisaccharide levels ^36^. Previous molecular dynamics simulation results suggested that in addition to the main site R113 located at the constriction of the pore lumen, K147, T145, and T115 were the second sensing sites of the acetamido group on glycans ^36^. The experimental results confirmed that M113R/T115A remained M113R for acetamido sensing and eliminating the secondary sensing site, which helped reduce the high blockage of glycan. [Data not shown]. We synthesized two pairs of glycans with differences only in glycosidic bonds. The pair of disaccharides were Galβ1, 3GlcNAc (di-1) and Galβ1, 4GlcNAc (di-2) (fig. 1a). The pair of tetrasaccharides were Galβ1, 3GlcNAcβ1, 3Galβ1, 3GlcNAc (tetra-1) and Galβ1, 4GlcNAcβ1, 3Galβ1, 4GlcNAc (tetra -2) (fig. 1b). They are two representative regioisomers, which differ in the glycosidic linkage position, used as model analytes for nanopore detection in our research. The nanopore measurements were carried out in a 2 M NaCI, 10 mM HEPES, pH 7.5 buffer, and -40 mV bias was continually applied. Each glycan was added to the *trans side* with a final concentration of 50 μM when a stable nanopore was formed to record the corresponding current trace (fig.1c). Interestingly, we found there’s another event based on level 1, which we defined as level 2 when analyzing current traces of these isomers (fig.1d). Three parameters were extracted from level 2, including the amplitude (Δ*I*_2_), dwell time (τ), and standard deviations (Std), to depict a 3D scatter plot ^37^ to identify the isomers (fig.1e).

**Figure 1.**
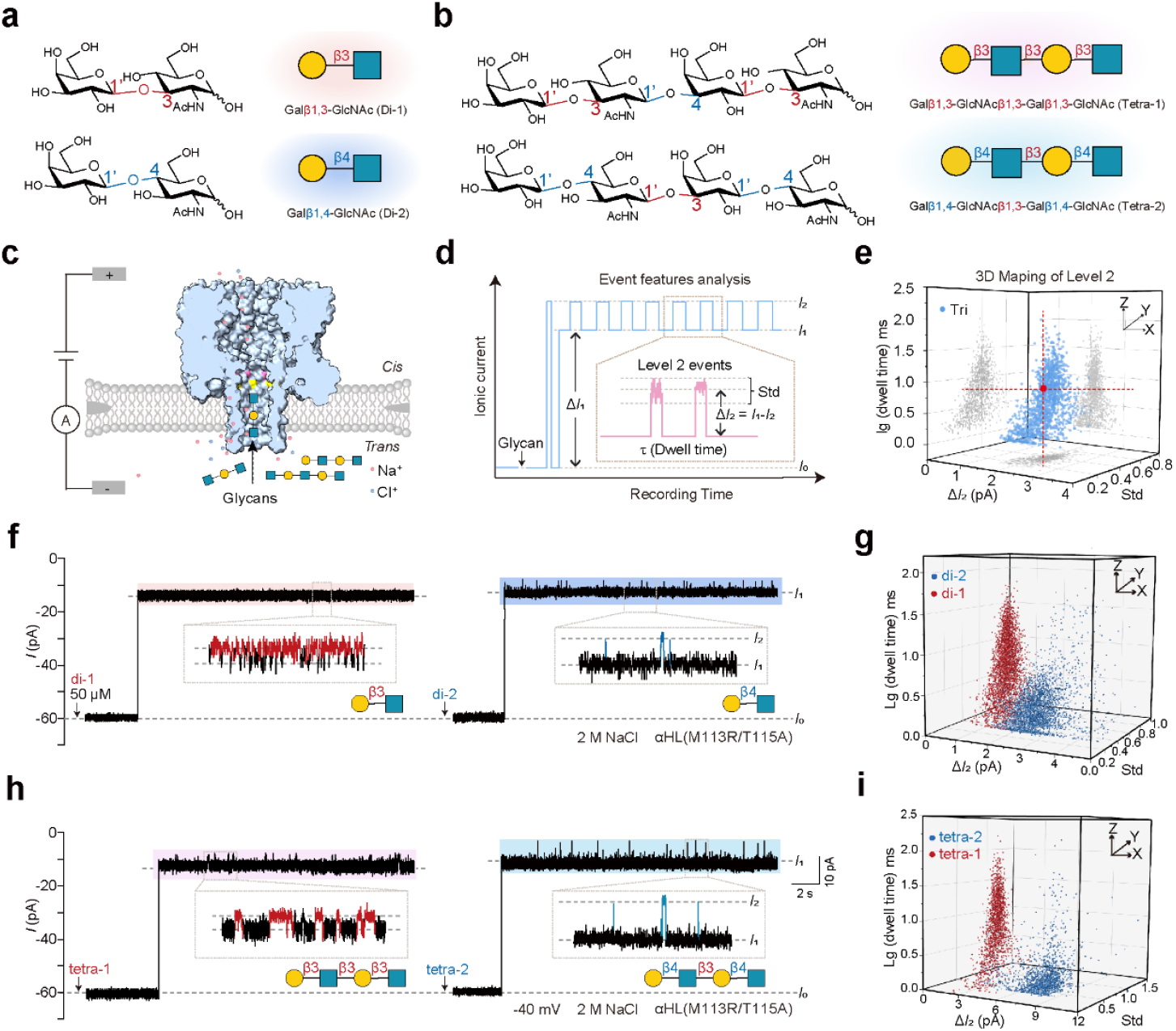
Differentiating glycans with different glycosidic bonds. (a-b) Structure of saccharides differing in glycosidic bonds. (c) Sectional view of α-HL (M113R/T115A) (part of the stem) cartoon structure, the mutant sites M113R and T115A were marked in pink and yellow, respectively. (d-e) Principle of nanopore-based glycan separation. (f) Representative ionic current traces of di-1 and di-2. The black arrow symbolizes the addition of glycan, with a final concentration of 50 μM. (g) 3D scatter plot of Δ*I*_2_ versus Std versus lg (dwell time) for mapping the level 2 events of di-1 and di-1 from the trace in (f). (h) Representative ionic current trace of glycan tetra-1 and tetra-2. (i) The 3D scatter plot of Δ*I*_2_ vs. Std vs. lg (dwell time) of tetra-1 and tetra-2 from the trace in (h).

Each pair of glycans showed distinct form of level 2 events (fig.1f and 1h). The stacked level 2 events fingerprints of two pairs of glycan regioisomers were shown in fig.1g and fig.1i. The scatter plots of level 2 events of di-1 and tetra-1 which are linked by only β1,3 glycosidic bonds exhibited elongated and narrow (wider dwell time range and narrower Δ*I*_2_ distribution range) features. Comparatively, the di-2 and tetra-2 containing both β1,4 and β1,3 glycosidic bonds correspond to shorter and broader 3D scatter plots. The Δ*I*_2_ dimension of di-1 di-2, tetra-1, and tetra-2 were centered at 1.58 pA, 2.64 pA, 3.71 pA, and 8.39 pA, respectively (fig.S1). It was easily found the Δ*I*_2_ value of the di-2 and tetra-2 (containing both β1,3 and β1,4 glycosidic linkage) were larger than di-1 and tetra-1 (containing only β1,3 glycosidic linkage), which suggested β1,4 linkage caused larger level 2 amplitude than that of β1,3 linkage. Besides, the two pairs of regioisomers also differed markedly in dwell time of level 2 events, with the lg (dwell time) mean values of 0.81, 0.23, 0.96, 0.12 for di-1 di-2, tetra-1 and tetra-2 respectively (fig.S1). Di-1 and tatra-1 showed longer dwell time than tetra-1 and tetra-2, indicating the β1,3 glycosidic linkage resulted in longer duration in the nanopore. The Std mean value of di-1, di-2, tetra-1, and tetra-2 were 0.37, 0.30, 0.47, and 0.30 (fig.S1). Compared with Δ*I*_2_ and dwell time, Std contributed minimum to regioisomer separation. Considering that different glycosidic bonds may lead to different orientations of functional groups in these glycans, thereby altering glycan-nanopore interaction, the distinct ionic current signal can be partly explained. To sum up, our results supported that glycans with distinct glycosidic bonds could be identified by αHL(M113R/T115A) nanopore.

To further investigate how α-HL(M113R/T115A) discriminates glycan isomers with different glycosidic linkages, we constructed MD simulation system of M113R/T115A with Gal1,3-GlcNAc and Gal1,4-GlcNAc, respectively (fig.2). The two disaccharides differ in only glycosidic bond. Each glycan added to the *trans* side of α-HL(M113R/T115A) was randomly diffused into the nanopore. Two disaccharide molecules were observed to occupy the stem lumen of α-HL simultaneously. Interestingly, Gal1,3-GlcNAc and Gal1,4-GlcNAc moved along completely different paths in lumen from the *trans*-side entrance to position nearby R113R/A115A. Meanwhile, the minimal permeation path radius tended to be narrower. When two Gal1,3-GlcNAc or two Gal1,4-GlcNAc approached R113/A115, the pore path radius and the relative distance between two glycan molecules kept fluctuating in distinct patterns, which could explain the different forms of level 2 events of glycans with varying glycosidic linkages and even other structure difference. It was also noticed that, compared with movement of GlcNAc molecule through α-HL(M113R) described in our previous work, both Gal1,3-GlcNAc and Gal1,4-GlcNAc appeared over twice longer time of movement in α-HL(M113R/T115A), and Gal1,3-GlcNAc or Gal1,4-GlcNAc bonding position was farther away from M113 site. These differences help to account for the higher level 2 event occurrence probability of M113R/T115A than M113R.

**Figure 2.**
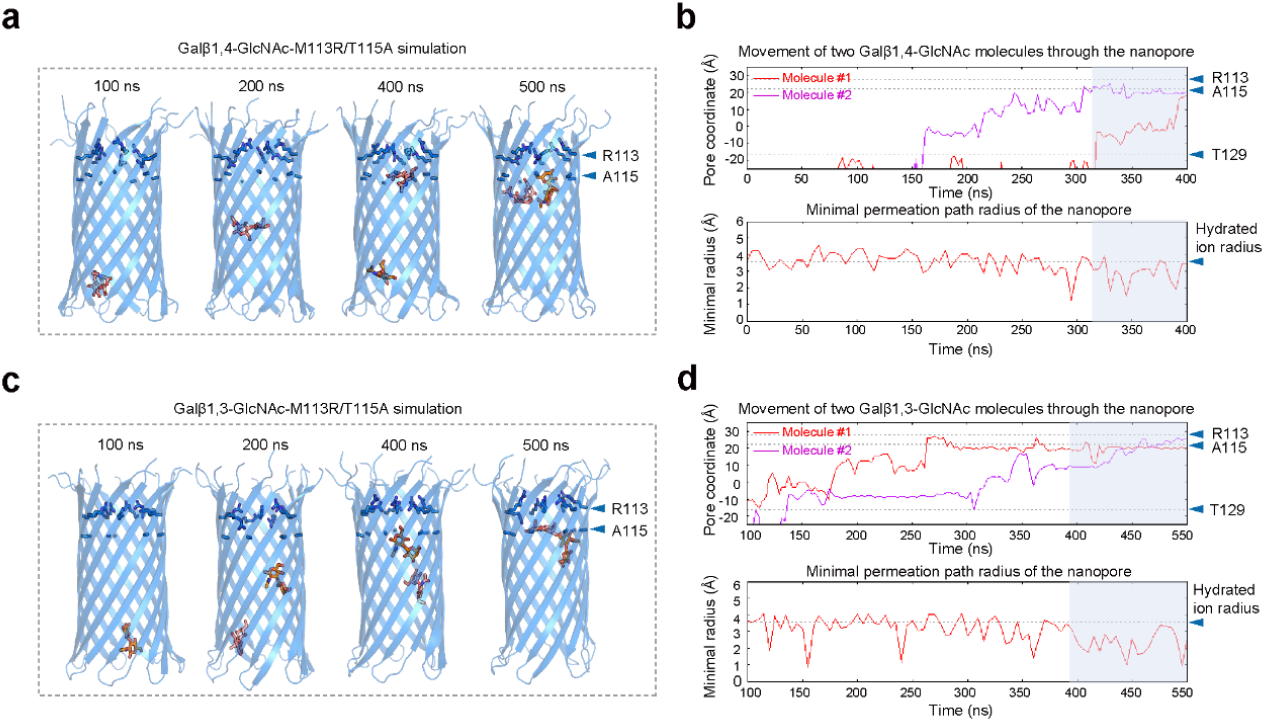
Molecular dynamics simulation for the behavior of glycans with different glycosidic bonds in the nanopore. (a) Movement of a Galβ1,4-GlcNAc-M113R/T115A molecule through nanopores in the simulation for Galβ1,4-GlcNAc - M113R/T115A system. (b) Galβ1,4-GlcNAc movement and permeation path radius as a function of time for Galβ1,4-GlcNAc - M113R/T115A system (c) Movement of a Galβ1,3-GlcNAc molecule through nanopores in simulation for Galβ1,3-GlcNAc - M113R/T115A system. (d) Galβ1,3-GlcNAc movement and permeation path radius as a function of time for the Galβ1,3-GlcNAc- M113R/T115A system.

### Distinguishment of glycans with different degrees of polymerization and conformation

Since saccharides differing in glycosidic bonds could be distinguished well, as shown in fig.1, it is worth exploring whether each of the saccharides can be discriminated when a mixture was added to the *trans side*. The identification strategy was depicted as shown in fig. 3a. In brief, di-1, di-2, tetra-1, and tetra-2 were mixed and then transferred to the *trans* side with a final concentration of 50 μM after a stable nanopore was formed on the lipid bilayer. Then the real-time signal of the mixture sensed by αHL(M113R/T115A) nanopore was recorded continuously for 1h. Level 2 events with distinct forms were observed during the recording (fig. 3b). Over 20000 Level 2 events were collected and their features including Δ*I*_2_ (amplitude of level2), dwell time, and Std (standard deviation) were extracted to draw the 2D ΔI_2_ versus lg (dwell time) or Std scatter plot of glycan mixture (fig.3c and fig.3d). The scatter plot can be divided into four groups exactly corresponding to level 2 events of four glycans. Both two kinds of 2D scatter of glycan mixture clustered mainly depending on Δ*I*_2_ dimension with the mean value of 1.52 pA, 2.44 pA, 4.24 pA, 8.37 pA. Lastly, the rapid identification of glycans was achieved by scatter projection of the mixture to the individual glycan. The distribution of four scatter groups of mixture glycan was consistent with the fingerprints of four glycans. The separation (s) values of Δ*I*_2_ Guassian fit for the pair of disaccharides and tetrasaccharides were calculated as, 90.20% and 400.08% respectively ^37, 38^. Compared to that of the disaccharides, the separation value of the tetrasaccharides was larger, possibly because the larger number difference of distinct glycosidic bonds in tetrasaccharides leads to greater changes in saccharides conformation, thus inducing larger ionic current difference. To our knowledge, although glycosidic linkages can be roughly determined by nuclear magnetic resonance (NMR) analysis, it becomes increasingly difficult for NMR as the length of glycan increases. By contrast, our results showed a better separation for tetrasaccharide than disaccharide using α-HL(M113R/T115A). Besides, the Δ*I*_*2*_ separation (s) values for di-1 and tetra-1 were calculated as 300.03%, and that for di-2 and tetra-2 was 400.27%, hinting the capability of M113R/T115A to distinguish different polymerization degrees. To date, the discrimination of distinct glycosidic bonds by traditional techniques (such as NMR, MS) showed shortcomings of indistinguishable differences ^20, 39^. Recently, a few researchers reported glycan isomer discrimination via nanopore depending on the chemical tag and machine learning ^34, 40^. Here, we provided a tag-free and high-resolution method for discrimination of both glycosidic linkage and polymerization degree that has never been reported to our knowledge.

**Figure 3.**
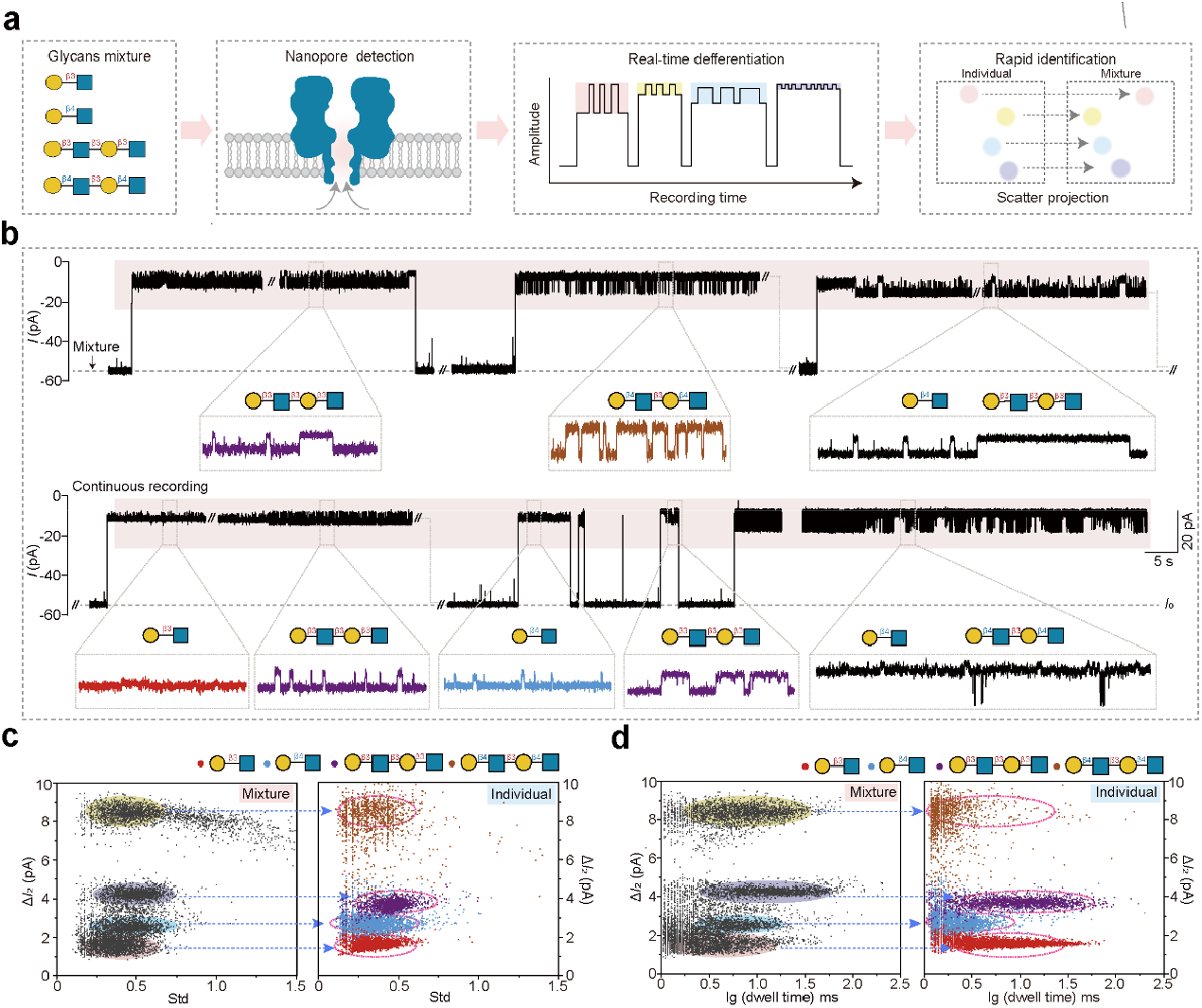
Identification of Glycans isomers with varying lengths in the mixture. (a) Workflow of glycan identification in the mixture. Step 1, mix four glycans with varying lengths and different glycosidic bonds at 50 μM final concentration. Step 2, αHL(M113R/T115A) nanopore sensing for the glycan mixture at the *trans* side. Step 3 (right), collect real-time signal of the glycans mixture. Step 4, rapid identification of glycans by scatter projection. (b) Representative current trace of the glycans mixture (upper). The typical level 2 events in the trace were enlarged and shown below (bottom), which were identified as four glycans (middle) by scatter projection shown in (c) and (d). The trace was recorded continuously for 40 minutes. (c) Scatter projection of Std versus Δ*I*_2_ from the level 2 events of the glycans mixture. The mixture scatter plot is on the left, and the fingerprint of four individual glycans is on the right. (d) Scatter projection of lg (dwell time) versus Δ*I*_2_ from the level 2 events of the glycans mixture. All the measurements were performed in 2 M NaCI, 10 mM HEPES, pH 7.5 symmetric buffer, with -40 mV voltage at the *trans* side. Each glycan was added to *trans* side to reach 50 μM final concentration.

## CONCLUSION

To date, discrimination of distinct glycosidic bonds is still a hurdle using traditional techniques (such as NMR, and MS) due to indistinguishable differences ^20, 39^. For example, chemical shifts of related carbon/hydrogen atoms from α-glucan with different glycosidic linkages are almost the same (<0.1) so discrimination between these saccharides is hard to achieve ^20^. But we are glad to see that a few research works on glycosidic bond discrimination have been reported recently. *Minmin Li et*.*al* successfully distinguished 6’-SL from 3’-SL, and *Shanyu Zhang et*.*al* achieved discrimination of disaccharide isomers of distinct glycosidic linkages, which are meaningful efforts to promote the development of nanopore technology in glycan isomer discrimination, but depending on the chemical tag and machine learning ^34, 40^. Hopefully, we have provided a tag-free and high-resolution method for glycosidic linkage discrimination that has never been reported to our knowledge. It is more difficult for NMR to characterize glycosidic linkage at the tetrasaccharide level than the disaccharide level. Interestingly, we found that nanopore resolution for tetrasaccharide isomer of glycosidic linkage was higher than disaccharide isomer. There are still some limitations in this research, (1) The type of glycosidic bond is limited to β-linkage, so further analysis should expand to other linkages except for β-1,3 and β-1,4 to make a more systematic detection. (2) Resolution between signals of saccharides is highly dependent on the nanopore, so developing other α-HL mutants or other biological nanopores is necessary to achieve discrimination for various glycosidic linkages. As the development of nanopore technology, discrimination of glycans with similar structures, including different lengths, and small individual monosaccharide units, that need to be detected in great demand, will inevitably make it hard to analyze with only an individual nanopore. We attempt to establish multiple nanopore platforms for detecting more kinds of physiologically and pathologically relevant glycan isomers in future work, which might provide foundations for the application of biological nanopores in glycan sequencing.

### Notes

The authors declare no competing financial interest.

## Acknowledgments

Dr. Zhaobing Gao and Dr. Bingqing Xia are grateful to Yi-Tao Long’s group (Nanjing University, China) for the help on data analysis, instrumentation, and helpful suggestions on this study. We are grateful to the National Science Fund of Distinguished Young Scholars (Grant 81825021), Fund of Youth Innovation Promotion Association (Grants 2019285, 2021333, and 2022077), the National Natural Science Foundation of China (Grants 81773707, 31700732, 82341058), Shanghai Municipal Science and Technology Major Project, Fund of Shanghai Science and Technology Innovation Action Plan (Grant 20ZR1474200), Shanghai Rising-Star Program (Grant 22QA1411000), Natural Science Foundation of Shanghai (Grant 22ZR1474000), the National Key Laboratory Program of China (Grants LG202101-01-04 and LG202102-01-01), the National Key Research and Development Program of China (Grant 2020YFC0842000), and the Strategic Leading Science and Technology Projects of Chinese Academy of Sciences (Grant XDA12050308) for financial support.

## MATERIALS AND METHODS

### Materials

Saccharides were purchased or synthesized by Wen Lab. Chemical reagents are as follows: sodium chloride (Sigma-Aldrich, USA), 4-(2-hydroxyethyl)-1-piperazineethanesulfonic acid (HEPES, Sigma-Aldrich, USA), sodium hydroxide (Sinopharm Chemical Reagent Co., Ltd, China), 1,2-diphytanoyl-sn-glycero-3-phosphocholine (DPhPC, Avanti Polar Lipids, USA), chloroform (Sinopharm Chemical Reagent Co., Ltd, China), decane (Sigma-Aldrich, USA), sodium dodecyl sulfate (SDS, Meilunbio, China), PrimeSTAR^®^ Max DNA Polymerase (Takara Biomedical Technology (Beijing) Co., Ltd, China), agarose (Yeason, China), Tris (Meilunbio, China), acetic acid (Sinopharm Chemical Reagent Co., Ltd, China), ethylene dinitrile tetra-acetic acid (EDTA, Sigma-Aldrich, USA), ampicillin sodium salt (Yeason, China), yeast extraction (Thermo Fisher, USA), tryptone (Thermo Fisher, USA), agar (Sinopharm Chemical Reagent Co., Ltd, China), ethanol (Sinopharm Chemical Reagent Co., Ltd, China), isopropanol (Sinopharm Chemical Reagent Co., Ltd, China), E.Z.N.A.® Plasmid DNA Mini Kit I (Omega Bio-Tek, China), isopropyl-*β*-D-1-thiogalactopyranoside (IPTG, Yeason, China), dithiothreitol (DTT, Yeason, China), Tween-20 (Sinopharm Chemical Reagent Co., Ltd, China), HisSep Ni-NTA agarose resin 6FF (Yeason, China), imidazole (Yeason, China), PageRuler™ Prestained Protein Ladder (Thermo Fisher, USA), Trans8K DNA marker (TransGen Biotech, China), SDS−PAGE electrophoresis buffer powder(Yeason, China), HRP-conjugated His-tag (Yeason, China), Peroxidase AffiniPure Goat Anti-Mouse IgG(H+L) (Yeason, China), Super ECL Detection Reagent (Yeason, China), Coomassie brilliant blue G250 (Yeason, China);

### Methods

#### Construction of Plasmids

The gene coding for α-hemolysin (wide type) was synthesized by Beijing Genomics Institute (BGI, China) and then inserted into the pEASY vector with a 6×histidine (His) tag at the C-terminus. For site-directed mutagenesis, the primers were designed on www.agilent.com.cn/store/primerDesignProgram.jsp. The plasmid of α-hemolysin (M113R) was constructed via polymerase chain reaction (PCR) with α-hemolysin (WT)-pEASY vector as the template (PCR program: 98 °C for 15 s; 55 °C for 15 s; 72 °C for 210 s for 30 cycles), using PrimeSTAR^®^ Max DNA Polymerase. The production of the mutant was firstly checked by gel electrophoresis, after which the product was treated with 1 μl Dpn I to remove the methylated template. Subsequently, the *E*.*coli trans5α* competent cell was transformed with the product via heat-shock and grown on an LB agar plate containing 100 μg/ml ampicillin. A single colony was picked for DNA sequencing to verify its sequence. The verified mutant was used for plasmid extraction by miniprep kit. α-hemolysin(M113R/T115A) was constructed similarly based on mutant M113R. All plasmids were verified for their genes before being used for nanopore purification.

#### Purification of Nanopores

In brief, the verified plasmids were transformed into the *E*.*coli BL21 pLysS* competent cells, and the *E*.*coli* strains were grown on LB agar plate, from which a single colony was chosen for DNA sequencing. The colony containing the correct plasmids was originally amplified in 30 ml LB medium with 100 μg/ml ampicillin at 37 °C/ 220 rpm for 14∼16 hours, then the cell suspension was transferred into 1.5 L LB medium with 100 μg/ml ampicillin and continued to grow at 37 °C/220 rpm. When OD_600_ of the suspension reached 0.6∼0.8, nanopore expression was induced with 1 mM IPTG at 22 °C / 200 rpm for 14∼18 hours. Afterward, the cell suspension was centrifuged at 4 °C/ 4000 rpm for 20 mins to collect the pellet, which was resuspended with TBST buffer containing 1 mM DTT and lysed by high-pressure homogenizer under 800−900 bar at 4 °C for 3∼5 mins. The soluble supernatant after centrifugation at 4 °C/ 13000 rpm for 30 mins was filtered with a 0.22 μm millipore. Then 1 mL Ni-NTA resin pre-equilibrated with TBS buffer was added to the supernatant and shaken at 4 °C for 1 hour to allow a sufficient binding to the protein. Subsequently, the mixture was transferred to a column and washed with linear gradient elution by imidazole (10 mM, 30 mM, 300 mM). After the protein was eluted with 300 mM imidazole, the solution was concentrated using Millipore (10KDa). Samples from each step were analyzed by SDS-PAGE.

#### Single-channel Recordings

In short, lipid bilayers of 1,2-diphytanoyl-sn-glycerol-3-phosphocholine (DPhPC, Avanti Polar Lipids) were formed across an aperture 200 μm in diameter on the cup that separated the chamber into two compartments, the *cis* side (grounded) and the *trans* side. And 1 ml electrolyte solution (1/2 M NaCl, 5 mM HEPES, pH7.5) was added to both sides. Unless otherwise stated, the protein was added to the *cis* side while the analytes were added to the *trans* side, and the voltage applied from the *trans* side was -40 mv. When a stable nanopore was formed on the bilayers, the current traces were recorded with a clamp amplifier (BC535, Warner Instruments, USA) and digitalized with a digitizer (Digidata 1550 A1 digitizer, Molecular Devices, U.K.) after the analytes were added to the *trans* side. The ionic current signals were low-pass filtered at 500 Hz and sampled at 10 kHz.

#### Data Analysis

Data analysis was performed with pClamp 10.2 software (Molecular Devices, USA). In general, the ignore duration set for level 1 and level 2 was 0.13 ms, 1ms respectively. The blockage ratio, defined as ΔI_1_/I_0_, was plotted by using Gaussian fit to a histogram of ΔI_1_/I_0_ values in GraphPad Prism 9 (Insightful Science, USA). A scatter plot of Δ*I*_2_ versus lg (Dwell time) or Std was also obtained using GraphPad Prism 9. The separation (*S*) between two analytes was then calculated by the eq below ^37^.

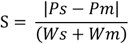

Where *Ps* and *Pm* represent for the peak values from the Gaussian fit of each analyte in a mixture plot and *Ws* and *Wm* are the corresponding width at half peaks of the Gaussian fit.

#### Molecular Dynamics Simulations

The nanopore models of M113R/T115A were built using X-ray structures of alpha-hemolysin as templates (PDB codes: 3M2L, 3M3R, 3M4D, 3M4E, 6U2S, 6U33, 6U3F, 6U3T, 6U49, 6U4P, 7AHL) ^35, 44, 45^. A modeller was used to generate the nanopore models ^46^. To build a simulation system, we placed the transmembrane domain of the nanopore into a 1-palmitoyl-2-oleoyl-sn-glycero-3-phosphocholine lipid bilayer. The lipid-embedded nanopore model was solvated in a periodic boundary condition box (120 × 120 × 185 A0) filled with TI3P water molecules and 0.15 M NaCl using CHARMM-GUI ^47^. Twenty-eight sugar molecules were randomly added to *trans*. The structure, topology, and parameter files of sugar molecules were generated using Glycan Modeler ^48^. Each system was replicated to perform three independent simulations. Based on the CHARMM36m all-atom force field ^49^, MD simulations were conducted using GROMACS 5.1.4 ^50^. After 100-ns equilibration, the nanopore models built based on a crystal structure of alpha-hemolysin (PDB code: 3M3R) produced stable conformations. Thus, a 1-μs production run was carried out for each simulation of these models. All productions were conducted in the isothermal–isobaric ensemble at a temperature of 303.15 K and a pressure of 1 atm. Temperature and pressure were controlled using the velocity-rescale thermostat ^51^ and the Parrinello-Rahman barostat with isotropic coupling ^52^, respectively. Equations of motion were integrated with a 2-fs time step, the LINCS algorithm was used to constrain bond length ^53^. Nonbonded pair lists were generated every 10 steps using a distance cutoff of 1.4 nm. A cutoff of 1.2 nm was used for Lennard-Jones (excluding scales 1 to 4) interactions, which were smoothly switched off between 1 and 1.2 nm. Electrostatic interactions were computed using a particle-mesh-Ewald algorithm with a real-space cutoff of 1.2 nm ^54^. The pore radius and ion-permeation path radius were calculated using HOLE ^55^.

## FIGURE LEGENDS

**Figure S1.**
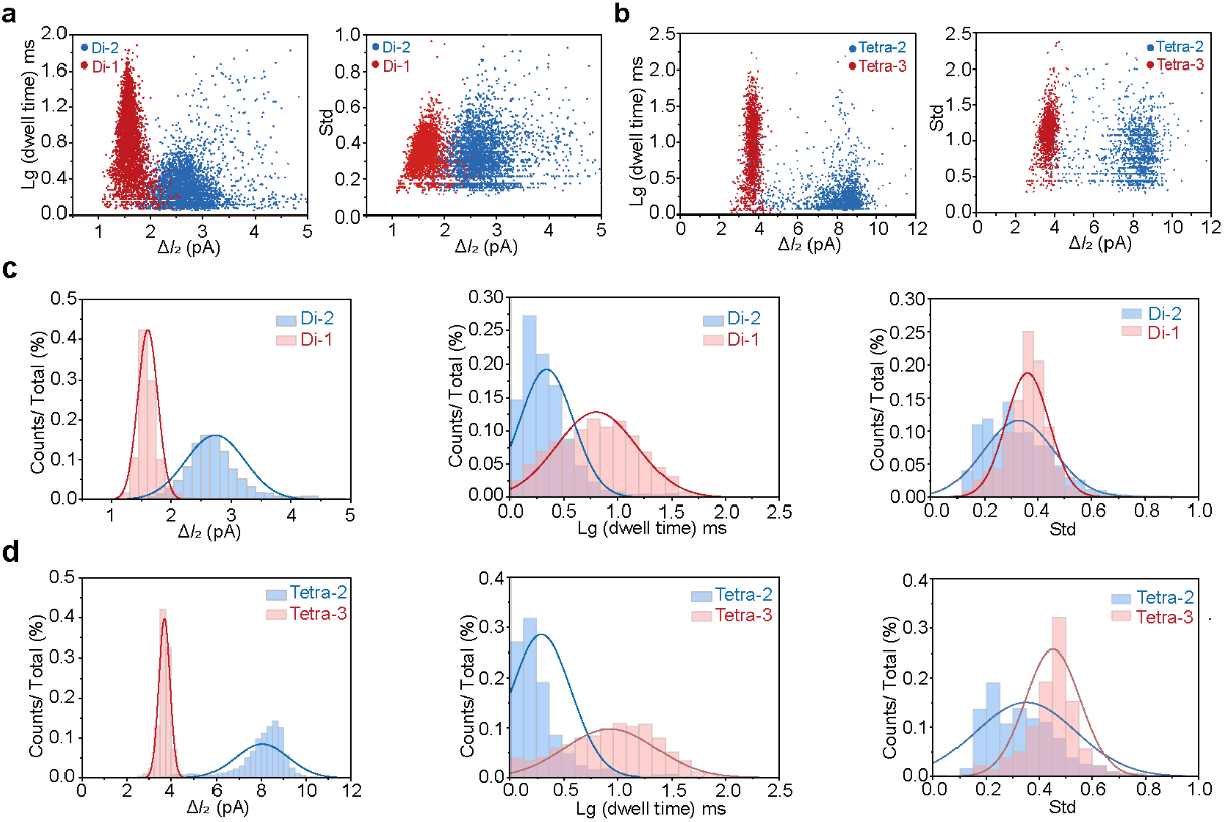
Level 2 events of glycans with different glycosidic linkages. (a) Δ*I*_2_ versus lg (dwell time) 2D scatter plot (Left) and Δ*I*_2_ versus Std 2D scatter plot (Right) of Di-1 and Di-2. (b) Δ*I*_2_ versus lg (dwell time) 2D scatter plot (Left) and Δ*I*_2_ versus Std 2D scatter plot (Right) of Tetra-2 and Tetra-3. (c) Histogram of Δ*I*_2_ (Left), lg (dwell time) (middle), and Std (Right) of Di-1 and Di-2 with Gaussian fit. (d) Histogram of Δ*I*_2_ (Left), lg (dwell time) (middle), and Std (Right) of Tetra-2 and Tetra-3 with Gaussian fit.

## REFERENCES

(1) Iozzo, R. V.; Schaefer, L. Proteoglycan form and function: A comprehensive nomenclature of proteoglycans. Matrix Biol 2015, 42, 11–55. DOI: 10.1016/j.matbio.2015.02.003.

(2) Kirschbaum, C.; Greis, K.; Mucha, E.; Kain, L.; Deng, S.; Zappe, A.; Gewinner, S.; Schöllkopf, W.; von Helden, G.; Meijer, G.; et al. Unravelling the structural complexity of glycolipids with cryogenic infrared spectroscopy. Nat Commun 2021, 12 (1), 1201. DOI: 10.1038/s41467-021-21480-1 From NLM.

(3) Flynn, R. A.; Pedram, K.; Malaker, S. A.; Batista, P. J.; Smith, B. A. H.; Johnson, A. G.; George, B. M.; Majzoub, K.; Villalta, P. W.; Carette, J. E.; et al. Small RNAs are modified with N-glycans and displayed on the surface of living cells. Cell 2021, 184 (12), 3109–3124.e3122. DOI: 10.1016/j.cell.2021.04.023 From NLM.

(4) Reily, C.; Stewart, T. J.; Renfrow, M. B.; Novak, J. Glycosylation in health and disease. Nat Rev Nephrol 2019, 15 (6), 346–366. DOI: 10.1038/s41581-019-0129-4 From NLM.

(5) Zeng, Y.; Xiang, Y.; Sheng, R.; Tomás, H.; Rodrigues, J.; Gu, Z.; Zhang, H.; Gong, Q.; Luo, K. Polysaccharide-based nanomedicines for cancer immunotherapy: A review. Bioact Mater 2021, 6 (10), 3358–3382. DOI: 10.1016/j.bioactmat.2021.03.008 From NLM.

(6) An, H. J.; Kronewitter, S. R.; de Leoz, M. L.; Lebrilla, C. B. Glycomics and disease markers. Curr Opin Chem Biol 2009, 13 (5-6), 601–607. DOI: 10.1016/j.cbpa.2009.08.015 From NLM.

(7) Phang, R.; Lin, C. H. Synthesis of Type-I and Type-II LacNAc-Repeating Oligosaccharides as the Backbones of Tumor-Associated Lewis Antigens. Front Immunol 2022, 13, 858894. DOI: 10.3389/fimmu.2022.858894 From NLM.

(8) Liu, Y.; Huang, P.; Jiang, B.; Tan, M.; Morrow, A. L.; Jiang, X. Poly-LacNAc as an age-specific ligand for rotavirus P[11] in neonates and infants. PLoS One 2013, 8 (11), e78113. DOI: 10.1371/journal.pone.0078113 From NLM.

(9) Chen, X. Human Milk Oligosaccharides (HMOS): Structure, Function, and Enzyme-Catalyzed Synthesis. Adv Carbohydr Chem Biochem 2015, 72, 113–190. DOI: 10.1016/bs.accb.2015.08.002 From NLM.

(10) Singh, A.; Kett, W. C.; Severin, I. C.; Agyekum, I.; Duan, J.; Amster, I. J.; Proudfoot, A. E. I.; Coombe, D. R.; Woods, R. J. The Interaction of Heparin Tetrasaccharides with Chemokine CCL5 Is Modulated by Sulfation Pattern and pH. J Biol Chem 2015, 290 (25), 15421–15436. DOI: 10.1074/jbc.M115.655845 From NLM.

(11) Williamson, K. A.; Hamilton, A.; Reynolds, J. A.; Sipos, P.; Crocker, I.; Stringer, S. E.; Alexander, Y. M. Age-related impairment of endothelial progenitor cell migration correlates with structural alterations of heparan sulfate proteoglycans. Aging Cell 2013, 12 (1), 139–147. DOI: 10.1111/acel.12031 From NLM.

(12) Roach, P. J. Glycogen phosphorylation and Lafora disease. Mol Aspects Med 2015, 46, 78–84. DOI: 10.1016/j.mam.2015.08.003 From NLM.

(13) Feyzi, E.; Saldeen, T.; Larsson, E.; Lindahl, U.; Salmivirta, M. Age-dependent modulation of heparan sulfate structure and function. J Biol Chem 1998, 273 (22), 13395–13398. DOI: 10.1074/jbc.273.22.13395 From NLM.

(14) Monneau, Y.; Arenzana-Seisdedos, F.; Lortat-Jacob, H. The sweet spot: how GAGs help chemokines guide migrating cells. J Leukoc Biol 2016, 99 (6), 935–953. DOI: 10.1189/jlb.3MR0915-440R From NLM.

(15) Ye, J.; Xia, H.; Sun, N.; Liu, C.-C.; Sheng, A.; Chi, L.; Liu, X.-W.; Gu, G.; Wang, S.-Q.; Zhao, J.; et al. Reprogramming the enzymatic assembly line for site-specific fucosylation. Nature Catalysis 2019, 2 (6), 514–522. DOI: 10.1038/s41929-019-0281-z.

(16) In Essentials of Glycobiology, Varki, A., Cummings, R. D., Esko, J. D., Stanley, P., Hart, G. W., Aebi, M., Mohnen, D., Kinoshita, T., Packer, N. H., Prestegard, J. H., et al. Eds.; Cold Spring Harbor Laboratory Press Copyright © 2022 by the Consortium of Glycobiology Editors, La Jolla, California. Published by Cold Spring Harbor Laboratory Press, Cold Spring Harbor, New York. All rights reserved., 2022.

(17) Varki, A. Biological roles of glycans. Glycobiology 2017, 27 (1), 3–49. DOI: 10.1093/glycob/cww086 From NLM.

(18) Lakshminarayanan, A.; Richard, M.; Davis, B. G. Studying glycobiology at the single-molecule level. Nature Reviews Chemistry 2018, 2 (8), 148–159. DOI: 10.1038/s41570-018-0019-5.

(19) Gray, C. J.; Migas, L. G.; Barran, P. E.; Pagel, K.; Seeberger, P. H.; Eyers, C. E.; Boons, G. J.; Pohl, N. L. B.; Compagnon, I.; Widmalm, G.; et al. Advancing Solutions to the Carbohydrate Sequencing Challenge. J Am Chem Soc 2019, 141 (37), 14463–14479. DOI: 10.1021/jacs.9b06406 From NLM.

(20) Fontana, C.; Widmalm, G. Primary Structure of Glycans by NMR Spectroscopy. Chem Rev 2023, 123 (3), 1040–1102. DOI: 10.1021/acs.chemrev.2c00580 From NLM.

(21) Yao, H. Y.; Wang, J. Q.; Yin, J. Y.; Nie, S. P.; Xie, M. Y. A review of NMR analysis in polysaccharide structure and conformation: Progress, challenge and perspective. Food Res Int 2021, 143, 110290. DOI: 10.1016/j.foodres.2021.110290 From NLM.

(22) Hofmann, J.; Hahm, H. S.; Seeberger, P. H.; Pagel, K. Identification of carbohydrate anomers using ion mobility–mass spectrometry. Nature 2015, 526 (7572), 241–244. DOI: 10.1038/nature15388.

(23) Schindler, B.; Barnes, L.; Renois, G.; Gray, C.; Chambert, S.; Fort, S.; Flitsch, S.; Loison, C.; Allouche, A. R.; Compagnon, I. Anomeric memory of the glycosidic bond upon fragmentation and its consequences for carbohydrate sequencing. Nat Commun 2017, 8 (1), 973. DOI: 10.1038/s41467-017-01179-y From NLM.

(24) Kim, Y.; Hyun, J. Y.; Shin, I. Glycan microarrays from construction to applications. Chem Soc Rev 2022, 51 (19), 8276–8299. DOI: 10.1039/d2cs00452f From NLM.

(25) Im, J.; Biswas, S.; Liu, H.; Zhao, Y.; Sen, S.; Biswas, S.; Ashcroft, B.; Borges, C.; Wang, X.; Lindsay, S.; et al. Electronic singlemolecule identification of carbohydrate isomers by recognition tunnelling. Nat Commun 2016, 7, 13868. DOI: 10.1038/ncomms13868 From NLM.

(26) Deamer, D.; Akeson, M.; Branton, D. Three decades of nanopore sequencing. Nat Biotechnol 2016, 34 (5), 518–524. DOI: 10.1038/nbt.3423 From NLM.

(27) Hu, Z. L.; Huo, M. Z.; Ying, Y. L.; Long, Y. T. Biological Nanopore Approach for Single-Molecule Protein Sequencing. Angew Chem Int Ed Engl 2021, 60 (27), 14738–14749. DOI: 10.1002/anie.202013462 From NLM.

(28) Ying, Y. L.; Hu, Z. L.; Zhang, S.; Qing, Y.; Fragasso, A.; Maglia, G.; Meller, A.; Bayley, H.; Dekker, C.; Long, Y. T. Nanopore-based technologies beyond DNA sequencing. Nat Nanotechnol 2022, 17 (11), 1136–1146. DOI: 10.1038/s41565-022-01193-2 From NLM.

(29) Xue, L.; Yamazaki, H.; Ren, R.; Wanunu, M.; Ivanov, A. P.; Edel, J. B. Solid-state nanopore sensors.

(30) Wang, Y.; Zhao, Y.; Bollas, A.; Wang, Y.; Au, K. F. Nanopore sequencing technology, bioinformatics and applications. Nat Biotechnol 2021, 39 (11), 1348–1365. DOI: 10.1038/s41587-021-01108-x From NLM.

(31) Kasianowicz, J. J.; Brandin, E.; Branton, D.; Deamer, D. W. Characterization of individual polynucleotide molecules using a membrane channel. Proc Natl Acad Sci U S A 1996, 93 (24), 13770–13773. DOI: 10.1073/pnas.93.24.13770 From NLM.

(32) Zhang, S.; Cao, Z.; Fan, P.; Wang, Y.; Jia, W.; Wang, L.; Wang, K.; Liu, Y.; Du, X.; Hu, C.; et al. A Nanopore-Based Saccharide Sensor. Angew Chem Int Ed Engl 2022, 61 (33), e202203769. DOI: 10.1002/anie.202203769 From NLM.

(33) Bayat, P.; Rambaud, C.; Priem, B.; Bourderioux, M.; Bilong, M.; Poyer, S.; Pastoriza-Gallego, M.; Oukhaled, A.; Mathé, J.; Daniel, R. Comprehensive structural assignment of glycosaminoglycan oligo- and polysaccharides by protein nanopore. Nat Commun 2022, 13 (1), 5113. DOI: 10.1038/s41467-022-32800-4 From NLM.

(34) Li, M.; Xiong, Y.; Cao, Y.; Zhang, C.; Li, Y.; Ning, H.; Liu, F.; Zhou, H.; Li, X.; Ye, X.; et al. Identification of tagged glycans with a protein nanopore. Nat Commun 2023, 14 (1), 1737. DOI: 10.1038/s41467-023-37348-5 From NLM.

(35) Song, L.; Hobaugh, M. R.; Shustak, C.; Cheley, S.; Bayley, H.; Gouaux, J. E. Structure of staphylococcal alpha-hemolysin, a heptameric transmembrane pore. Science 1996, 274 (5294), 1859–1866. DOI: 10.1126/science.274.5294.1859 From NLM.

(36) Xia, B.; Fang, J.; Ma, S.; Ma, M.; Yao, G.; Li, T.; Cheng, X.; Wen, L.; Gao, Z. Mapping the Acetylamino and Carboxyl Groups on Glycans by Engineered α-Hemolysin Nanopores. J Am Chem Soc 2023, 145 (34), 18812–18824. DOI: 10.1021/jacs.3c03563 From NLM.

(37) Xin, K. L.; Hu, Z. L.; Liu, S. C.; Li, X. Y.; Li, J. G.; Niu, H.; Ying, Y. L.; Long, Y. T. 3D Blockage Mapping for Identifying Familial Point Mutations in Single Amyloid-β Peptides with a Nanopore. Angew Chem Int Ed Engl 2022, 61 (44), e202209970. DOI: 10.1002/anie.202209970 From NLM.

(38) Hu, Z. L.; Li, Z. Y.; Ying, Y. L.; Zhang, J.; Cao, C.; Long, Y. T.; Tian, H. Real-Time and Accurate Identification of Single Oligonucleotide Photoisomers via an Aerolysin Nanopore. Anal Chem 2018, 90 (7), 4268–4272. DOI: 10.1021/acs.analchem.8b00096 From NLM.

(39) Dell, A.; Morris, H. R. Glycoprotein structure determination by mass spectrometry. Science 2001, 291 (5512), 2351–2356. DOI: 10.1126/science.1058890 From NLM.

(40) Zhang, S.; Cao, Z.; Fan, P.; Sun, W.; Xiao, Y.; Zhang, P.; Wang, Y.; Huang, S. Discrimination of Disaccharide Isomers of Different Glycosidic Linkages Using a Modified MspA Nanopore. Angew Chem Int Ed Engl 2023, e202316766. DOI: 10.1002/anie.202316766 From NLM.

(41) Lau, K.; Thon, V.; Yu, H.; Ding, L.; Chen, Y.; Muthana, M. M.; Wong, D.; Huang, R.; Chen, X. Highly efficient chemoenzymatic synthesis of β1–4-linked galactosides with promiscuous bacterial β1–4-galactosyltransferases. Chemical Communications 2010, 46 (33), 6066–6068. DOI: 10.1039/c0cc01381a.

(42) Li, Y.; Xue, M.; Sheng, X.; Yu, H.; Zeng, J.; Thon, V.; Chen, Y.; Muthana, M. M.; Wang, P. G.; Chen, X. Donor substrate promiscuity of bacterial β1–3-N-acetylglucosaminyltransferases and acceptor substrate flexibility of β1–4-galactosyltransferases. Bioorganic & Medicinal Chemistry 2016, 24 (8), 1696–1705. DOI: 10.1016/j.bmc.2016.02.043.

(43) McArthur, J. B.; Yu, H.; Chen, X. A Bacterial β1–3-Galactosyltransferase Enables Multigram-Scale Synthesis of Human Milk Lacto-N-tetraose (LNT) and Its Fucosides. ACS Catalysis 2019, 9 (12), 10721–10726. DOI: 10.1021/acscatal.9b03990.

(44) Banerjee, A.; Mikhailova, E.; Cheley, S.; Gu, L. Q.; Montoya, M.; Nagaoka, Y.; Gouaux, E.; Bayley, H. Molecular bases of cyclodextrin adapter interactions with engineered protein nanopores. Proc Natl Acad Sci U S A 2010, 107 (18), 8165–8170. DOI: 10.1073/pnas.0914229107.

(45) Liu, J.; Kozhaya, L.; Torres, V. J.; Unutmaz, D.; Lu, M. Structure-based discovery of a small-molecule inhibitor of methicillin-resistant Staphylococcus aureus virulence. J Biol Chem 2020, 295 (18), 5944–5959. DOI: 10.1074/jbc.RA120.012697.

(46) Sali, A.; Blundell, T. L. Comparative protein modelling by satisfaction of spatial restraints. J Mol Biol 1993, 234 (3), 779–815. DOI: 10.1006/jmbi.1993.1626.

(47) Wu, E. L.; Cheng, X.; Jo, S.; Rui, H.; Song, K. C.; Davila-Contreras, E. M.; Qi, Y.; Lee, J.; Monje-Galvan, V.; Venable, R. M.; et al. CHARMM-GUI Membrane Builder toward realistic biological membrane simulations. J Comput Chem 2014, 35 (27), 1997–2004. DOI: 10.1002/jcc.23702.

(48) Park, S. J.; Lee, J.; Qi, Y. F.; Kern, N. R.; Lee, H. S.; Jo, S.; Joung, I.; Joo, K.; Lee, J.; Im, W. CHARMM-GUI Glycan Modeler for modeling and simulation of carbohydrates and glycoconjugates. Glycobiology 2019, 29 (4), 320–331. DOI: 10.1093/glycob/cwz003.

(49) Guvench, O.; Mallajosyula, S. S.; Raman, E. P.; Hatcher, E.; Vanommeslaeghe, K.; Foster, T. J.; Jamison, F. W., 2nd; Mackerell, A. D., Jr. CHARMM additive all-atom force field for carbohydrate derivatives and its utility in polysaccharide and carbohydrate-protein modeling. J Chem Theory Comput 2011, 7 (10), 3162–3180. DOI: 10.1021/ct200328p.

(50) Van Der Spoel, D.; Lindahl, E.; Hess, B.; Groenhof, G.; Mark, A. E.; Berendsen, H. J. GROMACS: fast, flexible, and free. J Comput Chem 2005, 26 (16), 1701–1718. DOI: 10.1002/jcc.20291.

(51) Bussi, G.; Donadio, D.; Parrinello, M. Canonical sampling through velocity rescaling. J Chem Phys 2007, 126 (1). DOI: Artn 014101 10.1063/1.2408420.

(52) Aoki, K. M.; Yonezawa, F. Constant-pressure molecular-dynamics simulations of the crystal-smectic transition in systems of soft parallel spherocylinders. Phys Rev A 1992, 46 (10), 6541–6549. DOI: 10.1103/physreva.46.6541.

(53) Hess, B. P-LINCS: A Parallel Linear Constraint Solver for Molecular Simulation. J Chem Theory Comput 2008, 4 (1), 116–122. DOI: 10.1021/ct700200b.

(54) Darden, T.; York, D.; Pedersen, L. Particle Mesh Ewald - an N.Log(N) Method for Ewald Sums in Large Systems. J Chem Phys 1993, 98 (12), 10089–10092. DOI: Doi 10.1063/1.464397.

(55) Smart, O. S.; Goodfellow, J. M.; Wallace, B. A. The pore dimensions of gramicidin A. Biophys J 1993, 65 (6), 2455–2460. DOI: 10.1016/S0006-3495(93)81293-1.

